## Supplementary information for "Single nucleus RNA sequencing reveals glial cell type-specific responses to ischemic stroke"

Daniel Bormann[1,2], Michael Knoflach[3,4], Emilia Poreba[5], Christian J. Riedl[6], Giulia Testa[6], Cyrille Orset[7,8], Anthony Levilly[7,8], Andreá Cottereau[7,8], Phillip Jauk[6,9], Simon Hametner[6], Bahar Golabi[5], Dragan Copic[1,2,10], Katharina Klas[1,2], Martin Direder[1,2,11], Hannes Kühtreiber[1,2], Melanie Salek[1,2], Stephanie zur Nedden[12], Gabriele Baier-Bitterlich[12], Stefan Kiechl[3,4], Carmen Haider[6], Verena Endmayr[6], Romana Höftberger[6], Hendrik J. Ankersmit[1,2]*, Michael Mildner[5]*

[1] Applied Immunology Laboratory, Department of Thoracic Surgery, Medical University of Vienna, 1090 Vienna, Austria
[2]Aposcience AG, 1200 Vienna, Austria
[3]Department of Neurology, Medical University of Innsbruck, Anichstraße 35, 6020 Innsbruck, Austria
[4] VASCage, Research Centre on Vascular Ageing and Stroke, 6020 Innsbruck, Austria
[5]Department of Dermatology, Medical University of Vienna, 1090 Vienna, Austria
[6]Division of Neuropathology and Neurochemistry, Department of Neurology, Medical University of Vienna, 1090 Vienna, Austria
[7]Normandie University, UNICAEN, ESR3P, INSERM UMR-S U1237, Physiopathology and Imaging of Neurological Disorders (PhIND), GIP Cyceron, Institut Blood and Brain @ Caen-Normandie (BB@C), Caen, France
[8]Department of Clinical Research, Caen-Normandie University Hospital, Caen, France
[9]Center for Medical Physics and Biomedical Engineering, Medical University of Vienna, 1090 Vienna, Austria
[10]Division of Nephrology and Dialysis, Department of Internal Medicine III, Medical University of Vienna, 1090 Vienna, Austria.
[11]Department of Orthopedics and Trauma Surgery, Medical University of Vienna, 1090 Vienna, Austria
[12]Institute of Neurobiochemistry, CCB-Biocenter, Medical University of Innsbruck, 6020 Innsbruck, Austria
*: contributed equally

Correspondence to:

Michael Mildner, PhD
Department of Dermatology, Medical University of Vienna
Lazarettgasse 14, 1090 Vienna, Austria

or
Hendrik Jan Ankersmit, MD, MBA
Applied Immunology Laboratory, Department of Thoracic Surgery, Medical University of Vienna,
Waehringer Guertel 18-20, 1090 Vienna, Austria.


### Table of contents

**Supplementary Table 1.** Antibodies and antibody labelling kits used in this study.

**Supplementary Table 2.** Methodological details regarding immunofluorescence stainings.

**Supplementary Table 3.** Allen Brain Atlas database in situ hybridization (ISH) studies referenced within this study.

**Supplementary Figure 1.** Standardized tissue sampling and MRI validation of cerebral infarction.

**Supplementary Figure 2.** snRNAseq quality control and nuclei distribution across samples.

**Supplementary Figure 3.** A detailed transcriptional analysis of myeloid cells enriched within infarcted brain tissue.

**Supplementary Figure 4.** Global distribution of MCAO induced differentially expressed genes (DEGs) across all major cell clusters.

**Supplementary Figure 5.** DEG signatures of neurons within infarcted brain tissue.

**Supplementary Figure 6.** Additional pre-processing steps and between group comparisons pertaining to Figure 2.

**Supplementary Figure 7.** Analysis of DEGs in conserved oligodendrocyte lineage sub clusters.

**Supplementary Figure 8.** Limited transcriptional overlap between stroke associated oligodendrocyte lineage cells and diseases associated oligodendrocytes (DAO).

**Supplementary Figure 9.** Additional pre-processing steps and between group comparisons pertaining to Figure 4.

**Supplementary Figure 10.** Transcriptional signatures of reactive astrocytes in stroke and other neuropathologies.

**Supplementary Figure 11.** Cell-cell communication analysis infers immuno-glial cross talk within infarcted brain tissue.

**Supplementary Figure 12.** Cell-cell communication analysis infers intra-glial cross talk within infarcted brain tissue.

**Supplementary Figure 13.** Abundant proliferating, CD44 positive myeloid cells are identified in the perilesional zone surrounding the infarcted area.

**Supplementary Figure 14.** Spatial association of osteopontin positive myeloid cells to CD44 positive cells in human infarcted cerebral tissue.

**Supplementary notes** – Detailed description of cluster annotation.

**Supplementary references**

| Antibody | Source | Identifier |
|---|---|---|
| **Primary Antibodies** | | |
| Rabbit polyclonal anti-NG2 | Merck Millipore (Burlington, MA, USA) | Cat# AB5320, RRID:AB_91789 |
| Rabbit anti-NG2 - Cy3® conjugated | Merck Millipore (Burlington, MA, USA) | Cat# AB5320C3, RRID:AB_11203295 |
| Rabbit anti-BrdU - FITC conjugated | BD Bioscience (Franklin Lakes, NJ, USA) | From kit: Cat# 558662 |
| Rabbit polyclonal anti-Ki67 | Abcam (Cambridge, UK) | Cat# ab15580, RRID:AB_443209 |
| Rabbit polyclonal anti-CD44 | Abcam (Cambridge, UK) | Cat# ab157107, RRID:AB_2847859 |
| Recombinant rabbit monoclonal anti-Iba1 | Abcam (Cambridge, UK) | Cat# ab178846, RRID:AB_2636859 |
| Chicken polyclonal anti- GFAP | Abcam (Cambridge, UK) | Cat# ab4674, RRID:AB_304558 |
| Rabbit polyclonal anti- Osteopontin | Abcam (Cambridge, UK) | Cat# ab63856, RRID:AB_1524127 |
| Mouse monoclonal IgG1 anti-Vimentin, Clone V9 | DAKO - Agilent Technologies (Santa Clara, CA, USA) | Cat# M0725, RRID:AB_10013485 |
| Rabbit recombinant monoclonal anti-IL33 | Abcam (Cambridge, UK) | Cat# ab207737, RRID:AB_2827630 |
| **Secondary Antibodies** | | |
| Goat Anti-Rabbit IgG H&L (Alexa Fluor® 488) | Abcam (Cambridge, UK) | Cat# ab150077, RRID:AB_2630356 |
| Goat anti-Mouse IgG1 Cross-Adsorbed Secondary Antibody (Alexa Fluor® 546) | Thermo Fisher Scientific (Waltham, MA, USA) | Cat# A-21123, RRID:AB_2535765 |
| Goat anti-Chicken IgY (H+L) Secondary Antibody (Alexa Fluor® 647) | Thermo Fisher Scientific (Waltham, MA, USA) | Cat# A-21449, RRID:AB_2535866 |
| **Antibody labelling kits** | | |
| FlexAble CoraLite® Plus 488 Antibody Labeling Kit for Rabbit IgG | Proteintech (Rosemont, IL, USA) | Cat#: KFA001 |
| FlexAble CoraLite® Plus 555 Antibody Labeling Kit for Rabbit IgG | Proteintech (Rosemont, IL, USA) | Cat# KFA002 |
| FlexAble CoraLite® Plus 647 Antibody Labeling Kit for Rabbit IgG | Proteintech (Rosemont, IL, USA) | Cat# KFA003 |

**Supplementary Table 1.** Antibodies and antibody labelling kits used in this study.

| Colocalized targets | HIER | Non conjugated primary antibodies | | Corresponding secondary antibodies | | Fluorophore conjugated antibodies | | |
|---|---|---|---|---|---|---|---|---|
| | | Antibody | Dilution | Antibody | Dilution | Antibody | Labelling kit used | Final working con-centration |
| NG2+VIM+Ki67 | pH9 | NG2 | 1:100 | Goat Anti-Rabbit (Alexa Fluor® 488) | 1:500 | Ki67 | FlexAble CoraLite® Plus 647 | 2 µg/ml |
| | | VIM | 1:200 | Goat anti-Mouse (Alexa Fluor® 546) | 1:500 | | | |
| NG2+IL33+Ki67 | pH9 | NG2 | 1:100 | Goat Anti-Rabbit (Alexa Fluor® 488) | 1:500 | IL33 | FlexAble CoraLite® Plus 555 | 1.7 µg/ml |
| | | | | | | Ki67 | FlexAble CoraLite® Plus 647 | 2 µg/ml |
| GFAP+CD44+VIM | pH6 | GFAP | 1:800 | Goat anti-Chicken (Alexa Fluor® 647) | 1:400 | | | |
| | | VIM | 1:200 | Goat anti-Mouse (Alexa Fluor® 546) | 1:500 | | | |
| | | CD44 | 1:500 | Goat Anti-Rabbit (Alexa Fluor® 488) | 1:500 | | | |
| NG2+CD44+Ki67 | pH9 | NG2 | 1:100 | Goat Anti-Rabbit (Alexa Fluor® 488) | 1:500 | CD44 | FlexAble CoraLite® Plus 555 | 1 µg/ml |
| | | | | | | Ki67 | FlexAble CoraLite® Plus 647 | 2 µg/ml |
| Iba1+OPN+CD44 | pH6 | Iba1 | 1:400 | Goat Anti-Rabbit (Alexa Fluor® 488) | 1:500 | OPN | FlexAble CoraLite® Plus 555 | 3.3 µg/ml |
| | | | | | | CD44 | FlexAble CoraLite® Plus 647 | 1 µg/ml |
| Iba1+CD44+Ki67 | pH6 | Iba1 | 1:400 | Goat Anti-Rabbit (Alexa Fluor® 488) | 1:500 | CD44 | FlexAble CoraLite® Plus 555 | 1 µg/ml |
| | | | | | | Ki67 | FlexAble CoraLite® Plus 647 | 2 µg/ml |

**Supplementary Table 2.** Methodological details regarding immunofluorescence stainings. Antibody combinations, heat induced epitope retrieval (HIER) pH and antibody dilutions are summarized for each performed immunofluorescence staining.

| Marker Gene | Main associated cell populations | Allen Atlas ISH Study | Hyperlink |
|---|---|---|---|
| Rasgrf2 | L2/L3 | Rasgrf2 - RP_051214_01_F08 - coronal | https://mouse.brain-map.org/gene/show/19181 |
| Rorb | L4/L5 | Rorb - RP_071018_01_H03 - coronal | https://mouse.brain-map.org/experiment/show/79556597 |
| Serpine2 | L5 | Serpine2 - RP_051214_01_H06 - coronal | https://mouse.brain-map.org/gene/show/20482 |
| Grik3 | L5/6 | Grik3 - RP_060606_03_A02 - coronal | https://mouse.brain-map.org/experiment/show/75749418 |
| Tle4 | L5/6 | Tle4 - RP_051101_03_A08 - coronal | https://mouse.brain-map.org/experiment/show/73521809 |
| Syt6 | L6 | Syt6 - RP_Baylor_103197 – coronal | https://mouse.brain-map.org/gene/show/33815 |
| Synpr | L6/CLA | Synpr - RP_040507_02_G03 - coronal | https://mouse.brain-map.org/gene/show/47844 |
| Gng2 | L6/CLA | Gng2 - RP_040604_01_F06 - coronal | https://mouse.brain-map.org/experiment/show/67936006 |
| Ndst4 | LSr, PIR | Ndst4 - RP_060220_03_B07 - coronal | https://mouse.brain-map.org/gene/show/41177 |

**Supplementary Table 3.** Allen Brain Atlas database *in situ* hybridization (ISH) studies [25] referenced within this study. Abbreviations: L = layer (cortical layers L2 –L6), CLA = claustrum, LSr = lateral septal nucleus, PIR = piriform cortex.

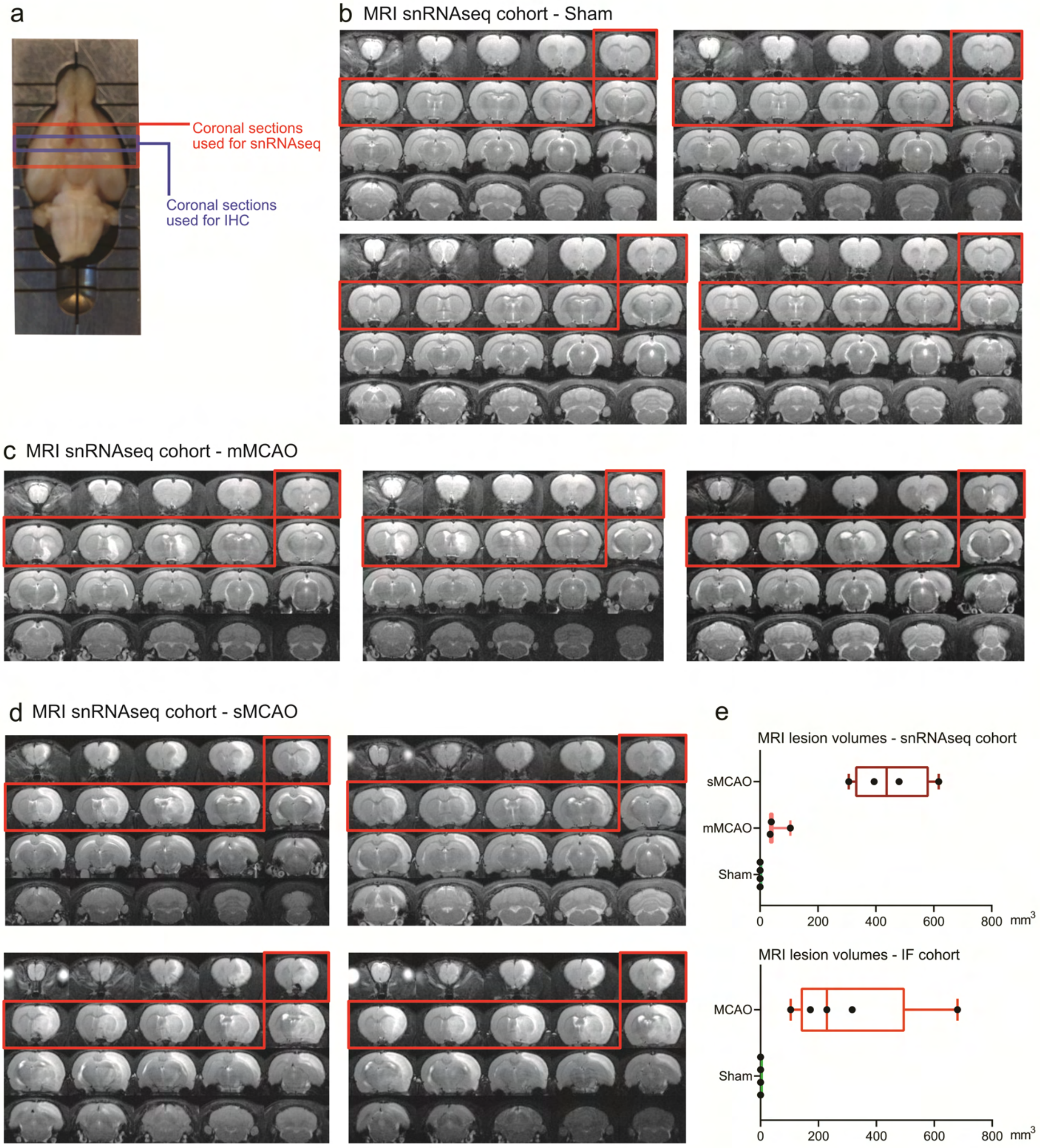


**Supplementary Figure 1. Standardized tissue sampling and MRI validation of cerebral infarction. (a)** Representative photograph of a rat brain placed in the adult rat brain slicer matrix, used for dissecting coronal brain sections. The red box denotes the brain region sampled for snRNAseq studies, the blue box depicts the brain region used for immunofluorescence staining. **(b-d)** T2-weighted MRI images of all animals used for snRNAseq in the Sham group (**b**), as well as MCAO group (**c** and **d**) are shown, hyper intense areas demark infarcted tissue. The MCAO group was further subdivided in cases with moderate (mMCAO) (**c**) and severe (sMCAO) infarctions (**d**). **(e)** Distributions of overall infarct lesion sizes in the rat cohorts used for snRNAseq (top) and immunofluorescence staining (bottom) are presented as boxplots, depicting medians, 25th to 75th percentiles as hinges, minimal and maximal values as whiskers, and individual lesion volumes for each animal as dots.

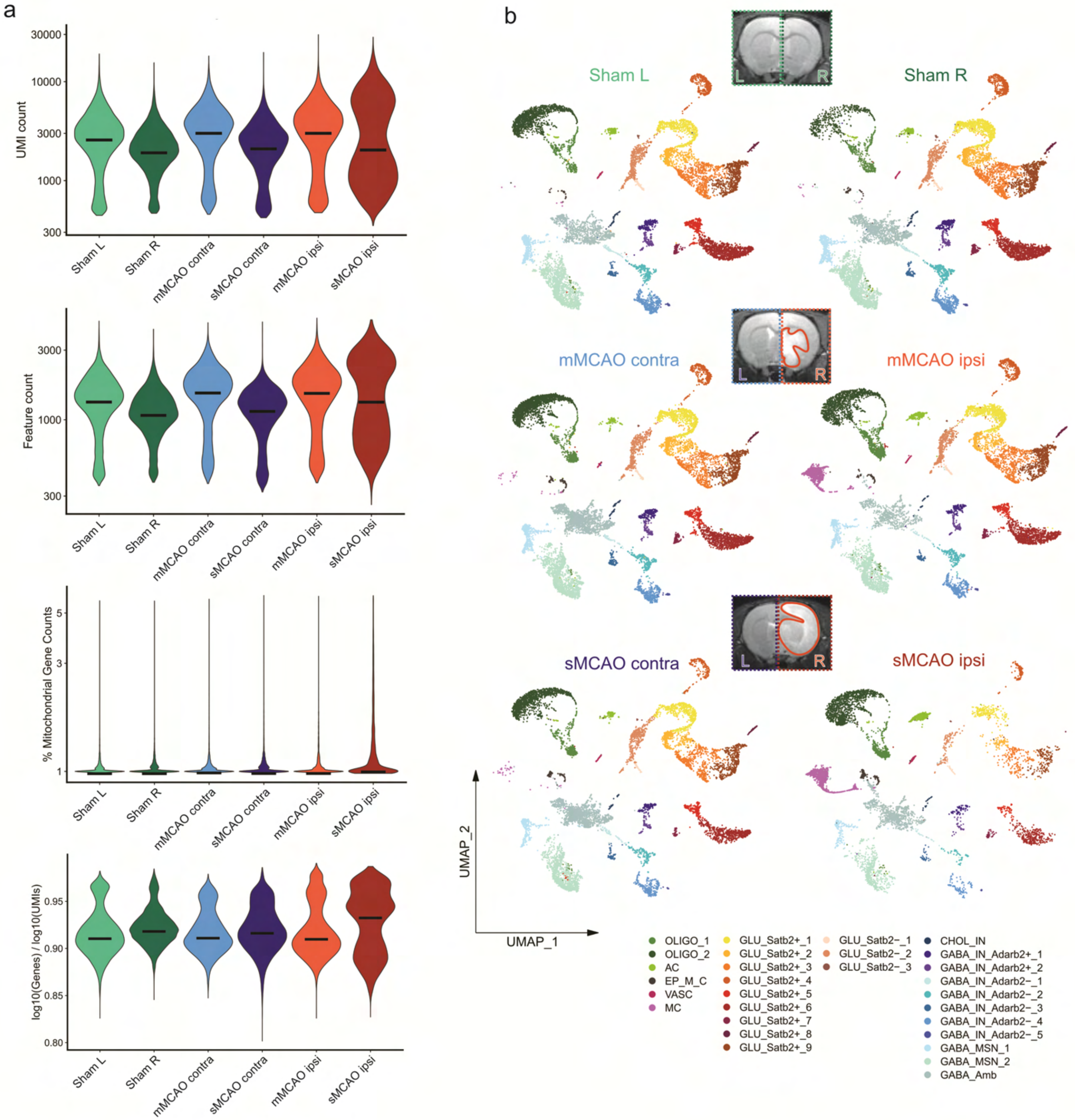


**Supplementary Figure 2. snRNAseq quality control and nuclei distribution across samples. a** Major quality control metrics derived from the final integrated dataset. From top to bottom: Number of unique transcripts (=UMIs), genes (=features), percentages of reads aligned to mitochondrial genes and the ratio of log10 of gene counts to log10 UMI counts (novelty score). All metrics are reported for each sequenced sample, with y axes scales log normalized. Black bars denote median values. **(b)** UMAP plots depict distribution of nuclei across major cell clusters, split by individual samples. T2-weighted MRI images show representative coronal brain sections of Sham group rats (top) and MCAO group rats with moderate (mMCAO) (middle) and severe infarctions (sMCAO) (bottom).

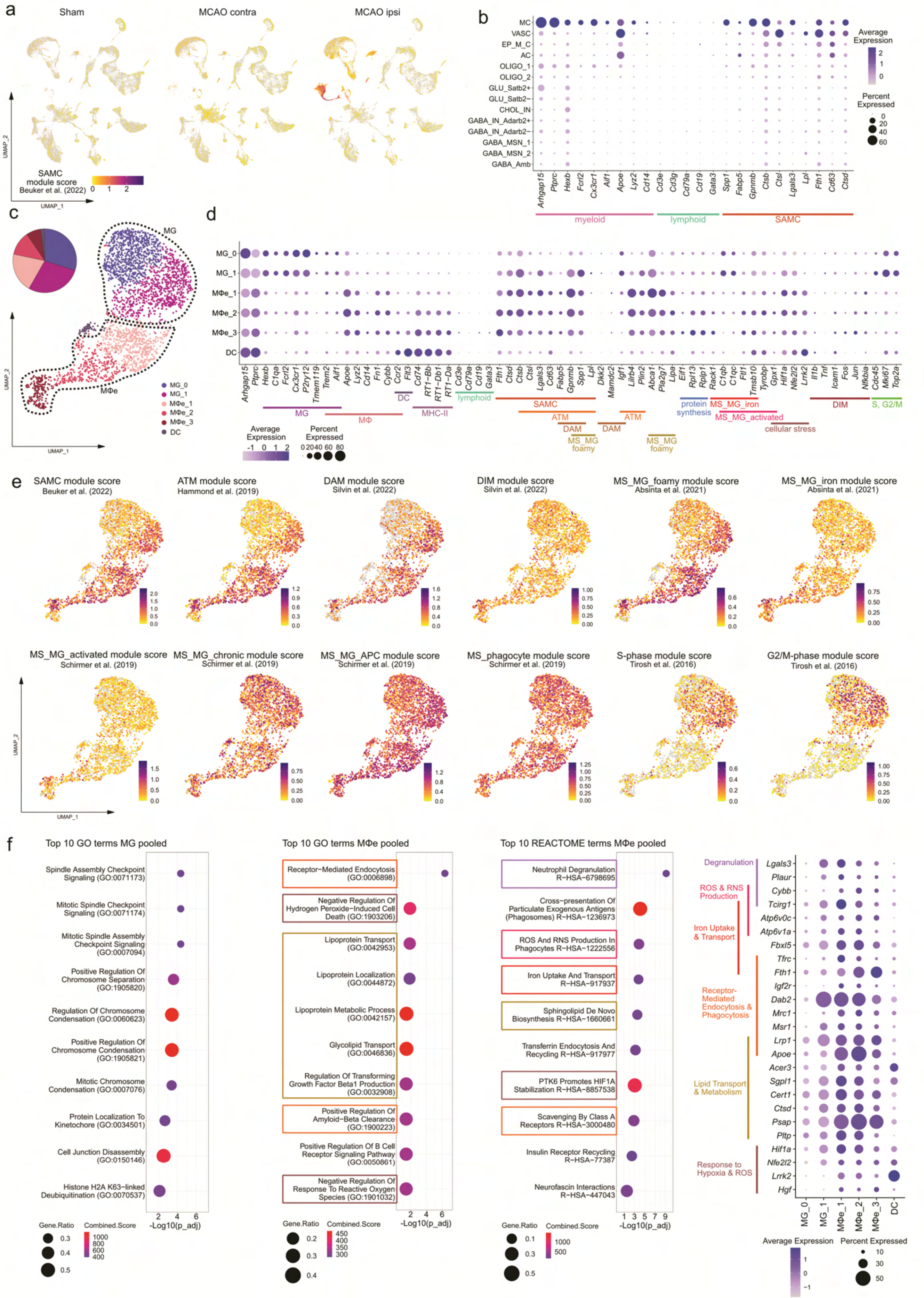


**Supplementary Figure 3. A detailed transcriptional analysis of myeloid cells enriched within infarcted brain tissue**. (Legend on next page)

**Supplementary Figure 3. A detailed transcriptional analysis of myeloid cells enriched within infarcted brain tissue**. **(a)** Feature plots showing stroke associated myeloid cell (SAMC) gene module score expression projected onto main clustering UMAP plots, split by treatment group. **(b)** Dotplot depicting expression of canonical myeloid and lymphoid cell lineage markers, as well as the genes making up the SAMC module gene set within the main clusters of the integrated dataset. **(c)** Subclustering of myeloid cells derived from MCAO ipsi. UMAP plot depicting 2646 nuclei annotated to 6 sub clusters. The relative contribution of each subcluster to all MCAO ipsi enriched myeloid cells is presented as pie plot. **(d)** Dotplot depicting the expression of canonical microglia, macrophage, DC, MHC-II and lymphoid cell associated genes, as well as representative genes from previously described MC gene sets within each myeloid subcluster. **(e)** Module score feature plots projecting aggregate expressions of various gene sets on to MC subcluster UMAP plots. Abbreviations: MG: microglia, MΦ: macrophage, MΦe: macrophage enriched, DC: dendritic cell, MHC: major histocompatibility complex, SAMC: stroke associated myeloid cell, ATM: Axon Tract-Associated Microglia, DAM: disease-associated microglia, MS_MG: Multiple sclerosis associated microglia, DIM: disease inflammatory macrophage, APC: antigen-presenting cell. **(f)** Results of functional enrichment analyses of MG and MΦe cluster marker genes (log2FC >= 0.6, Bonferroni-adjusted p-values < 0.05) are depicted as dot plots. Top 10 enriched GO biological process terms, subset by combined score and order by $-\log_{10}$ of Benjamini-Hochberg method adjusted p values are shown for each subset. Top10 enriched REACTOME terms were derived from MΦe cluster markers. Outer right dotplot depicts the average expression of representative genes, associated to the top enriched terms derived from MΦe cluster markers, for each MC subcluster. Color coded functional annotations are given next to gene names on the y-axis.

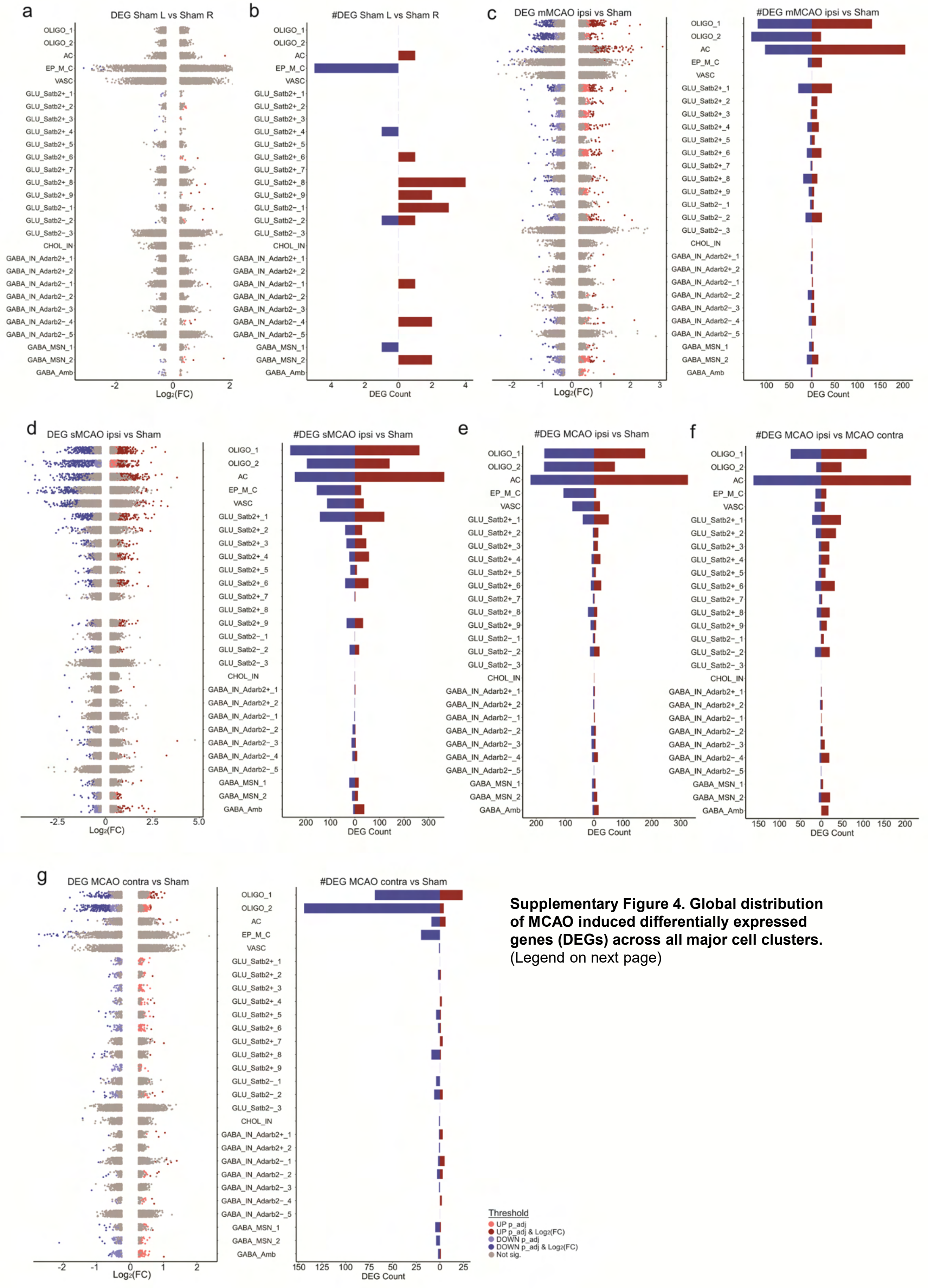


**Supplementary Figure 4. Global distribution of MCAO induced differentially expressed genes (DEGs) across all major cell clusters.** (Legend on next page)

**Supplementary Figure 4. Global distribution of MCAO induced differentially expressed genes (DEGs) across all major cell clusters.** Results of DEG calculations for each conserved cell cluster comparing various groups. Bar plots depict the numbers of up- and downregulated DEGs, meeting cut offs for statistical significance (Bonferroni-adjusted p-values < 0.05) and magnitude of change in gene expression (|log2fold change ≥ 0.6|) (=DEG count). Strip plots depict distribution of DEGs, color coded by DEG cut offs, depicted in the lower right legend within the figure. Results for the following comparisons are shown: **a,b** DEG distribution **(a)** and DEG counts **(b)** from the comparison of the left (L) and right (R) hemisphere datasets derived from Sham control animals. **c,d** DEG distribution left, DEG counts right from the comparison of datasets derived from hemispheres with moderate (mMCAO ipsi) (**c**) or severe (sMCAO ipsi) infarction (**d**) to Sham control datasets. **e,f** DEG counts derived from the comparison of MCAO infarcted hemisphere datasets (moderate and severe pooled) (MCAO ipsi) to Sham control datasets (Sham) (**e**) and datasets derived from the hemisphere contralateral to infarction (MCAO contra) (**f**), corresponding strip plots are shown in figure 1. **g** DEG distribution left, DEG counts right from the comparison of datasets derived from hemispheres contralateral to infarction (MCAO contra) to Sham datasets.

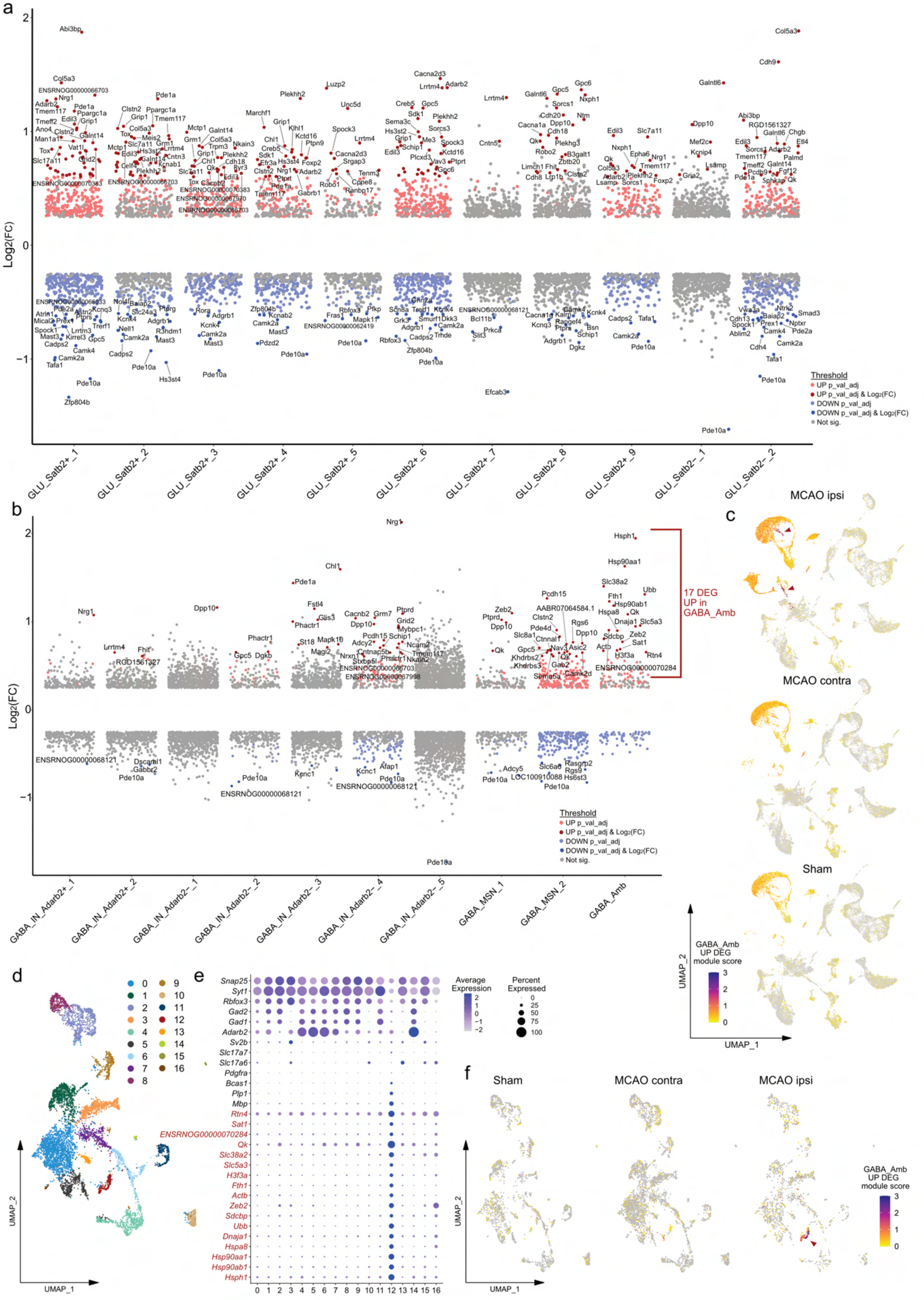


**Supplementary Figure 5. DEG signatures of neurons within infarcted brain tissue.** (Legend on next page)

**Supplementary Figure 5. DEG signatures of neurons within infarcted brain tissue. (a,b)** Results of DEG calculations for glutamatergic **(a)** and GABAergic **(b)** neuronal cell clusters, comparing gene expressions within datasets derived from MCAO ipsi to datasets from MCAO contra, for each neuronal cluster. DEGs are presented as color coded strip plots, up to top 20 significantly (adjusted p-values <0.05) up- and downregulated DEGs, sorted by log2FC are labelled. **(c)** Module score feature plots depicting the aggregate expression of the 17 DEGs upregulated in the ambiguous GABAergic neuronal cluster (GABA_Amb) in infarcted brain tissue. Feature plots are split by group, red arrowheads highlight UMAP regions with highest module scores, DEGs included in the module score are highlighted in **(b)**. **(d)** UMAP plot depicting unsupervised subclustering analysis of GABA_Amb. **(e)** Dotplot showing the expression of canonical pan neuronal, GABAergic and glutamatergic neuronal subset markers, as well as oligodendrocyte lineage associated markers and the 17 DEGs upregulated in the GABA_Amb cluster within MCAO ipsi across all GABA_Amb subclusters. **(f)** Module score feature plots projecting the aggregate expression of the 17 DEGs upregulated in GABA_Amb in infarcted brain tissue onto the GABA_Amb subclustering UMAP plot, split by treatment group.

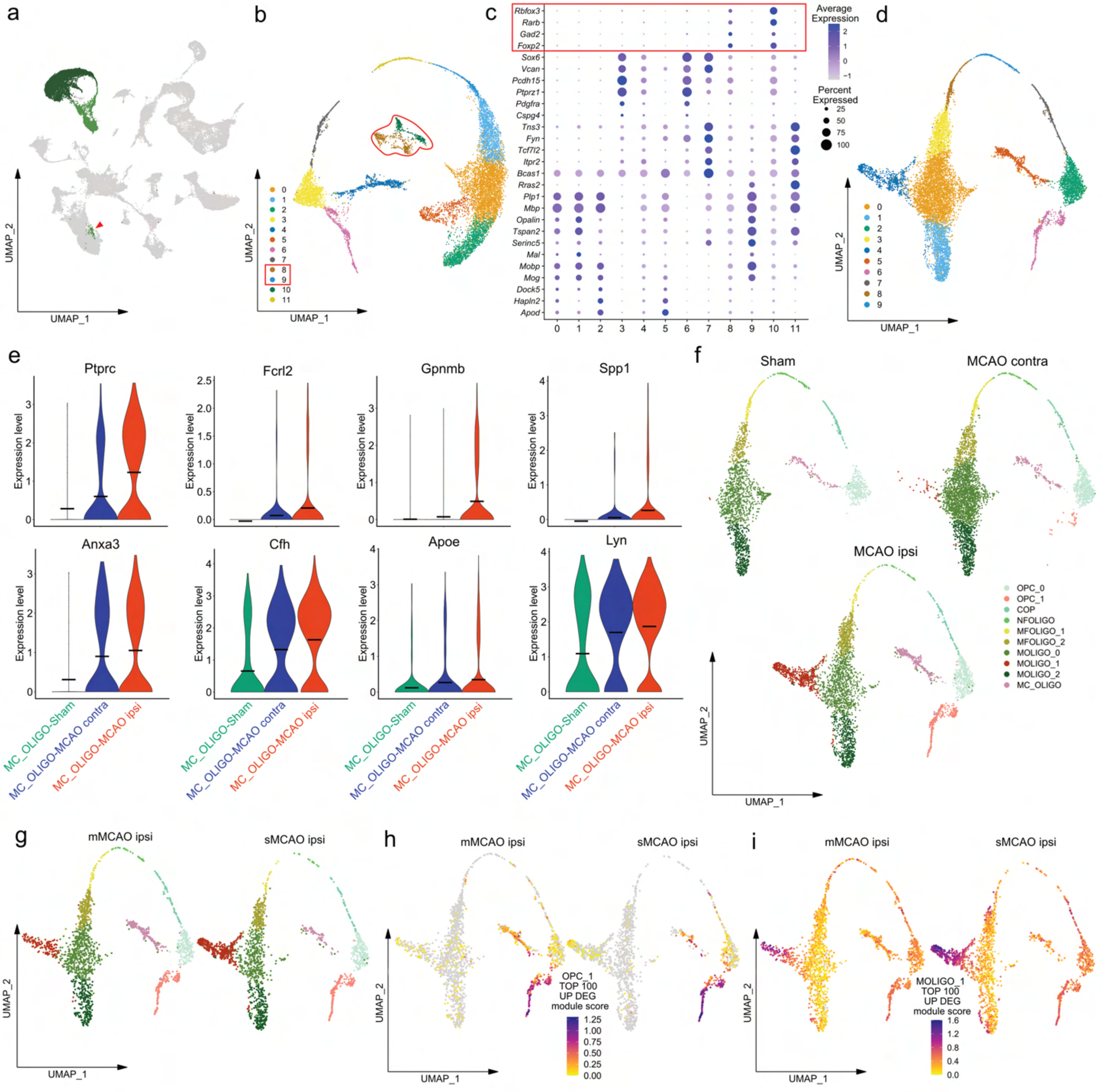


**Supplementary Figure 6. Additional pre-processing steps and between group comparisons pertaining to Figure 2. (a)** Oligodendrocyte lineage clusters OLIGO_1 and OLIGO_2 are highlighted in the main clustering UMAP plot (green). The red arrowhead points to scattered OLIGO_1 nuclei in proximity to the neuronal cluster MSN_2. **(b)** UMAP showing unsupervised subclustering of oligodendrocyte lineage clusters, depicting color coded Seurat clusters. **(c)** Dotplot depicting curated neuronal and oligodendrocyte marker genes. Clusters 8 and 9 and medium spiny neuron (MSN) signature genes specifically expressed within these cluster are highlighted in red. **(d)** UMAP plot depicting oligodendrocyte subclusters, after removal of contaminating subclusters. This subclustering was used for subsequent analyses. **(e)** Violin plots showing gene expression levels of myeloid cell associated genes within the myeloid cell oligodendrocyte mixed subcluster (MC_OLIGO), split by treatment group. **(f,g)** UMAP plots depict distribution of nuclei across subclusters, split by treatment group **(f)** and infarction severity **(g)**. **(h, i)** The top 100 significantly (Bonferroni-adjusted p-values < 0.05) upregulated DEGs, sorted by descending $\log_2$fold changes, derived from the comparison of subcluster OPC_1 to OPC_0 **(h)** and MOLIGO_1 to MOLIGO_2 **(i)** were aggregated to module scores and projected onto sub cluster UMAP plots, split by severity.

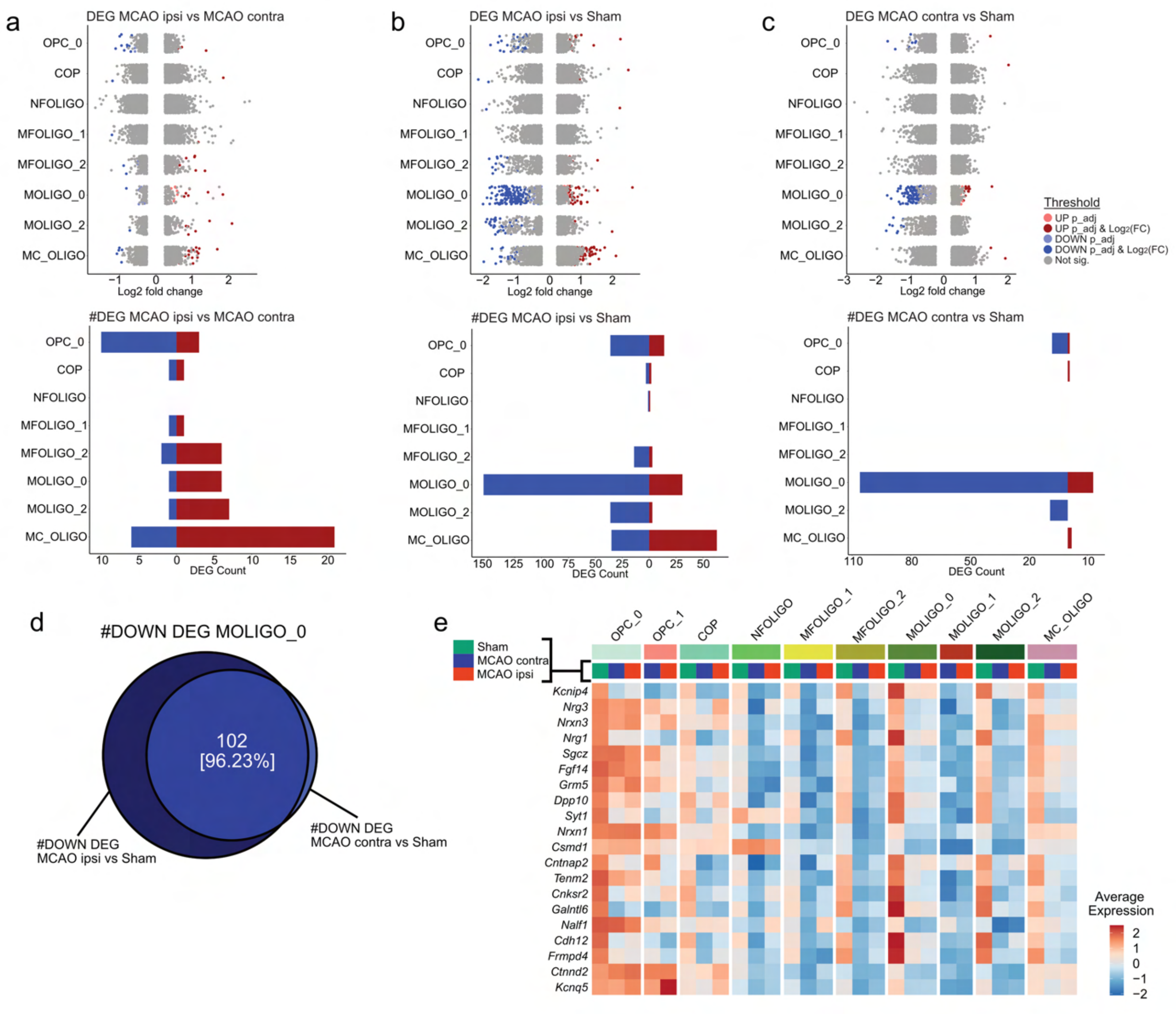


**Supplementary Figure 7. Analysis of DEGs in conserved oligodendrocyte lineage sub clusters.** Results of DEG calculations for each conserved oligodendrocyte lineage sub cluster, comparing various groups. **(a-c)** Strip plots (top) depicting distribution of DEGs, color coded by DEG cut offs. Bar plots (bottom) depict the numbers of up- and downregulated DEGs, meeting cut offs for statistical significance (Bonferroni-adjusted p-values < 0.05) and magnitude of change in gene expression (|log2fold change ≥ 0.6|) (= DEG count). Results for the following comparisons are shown: Datasets derived from MCAO ipsi compared to datasets derived from MCAO contra or Sham controls (**a** and **b**, respectively), **(c)** Datasets derived from MCAO contra vs Sham datasets. **(d)** Venn diagram depicting the overlap between downregulated DEGs within subcluster MOLIGO_0 in MCAO ipsi relative to Sham (left circle) and MCAO contra relative to Sham (right circle). Numbers denote the absolute and relative numbers of downregulated DEGs derived from the MCAO contra vs Sham comparison, which are also downregulated in MCAO ipsi relative to Sham datasets. **(e)** Heatmap depicting the average scaled gene expression of the Top 20 DEGs downregulated in MCAO contra relative to Sham, as sorted by $log_2$fold changes, split by sub cluster and group. All of these genes were also significantly downregulated in MCAO ipsi relative to Sham.

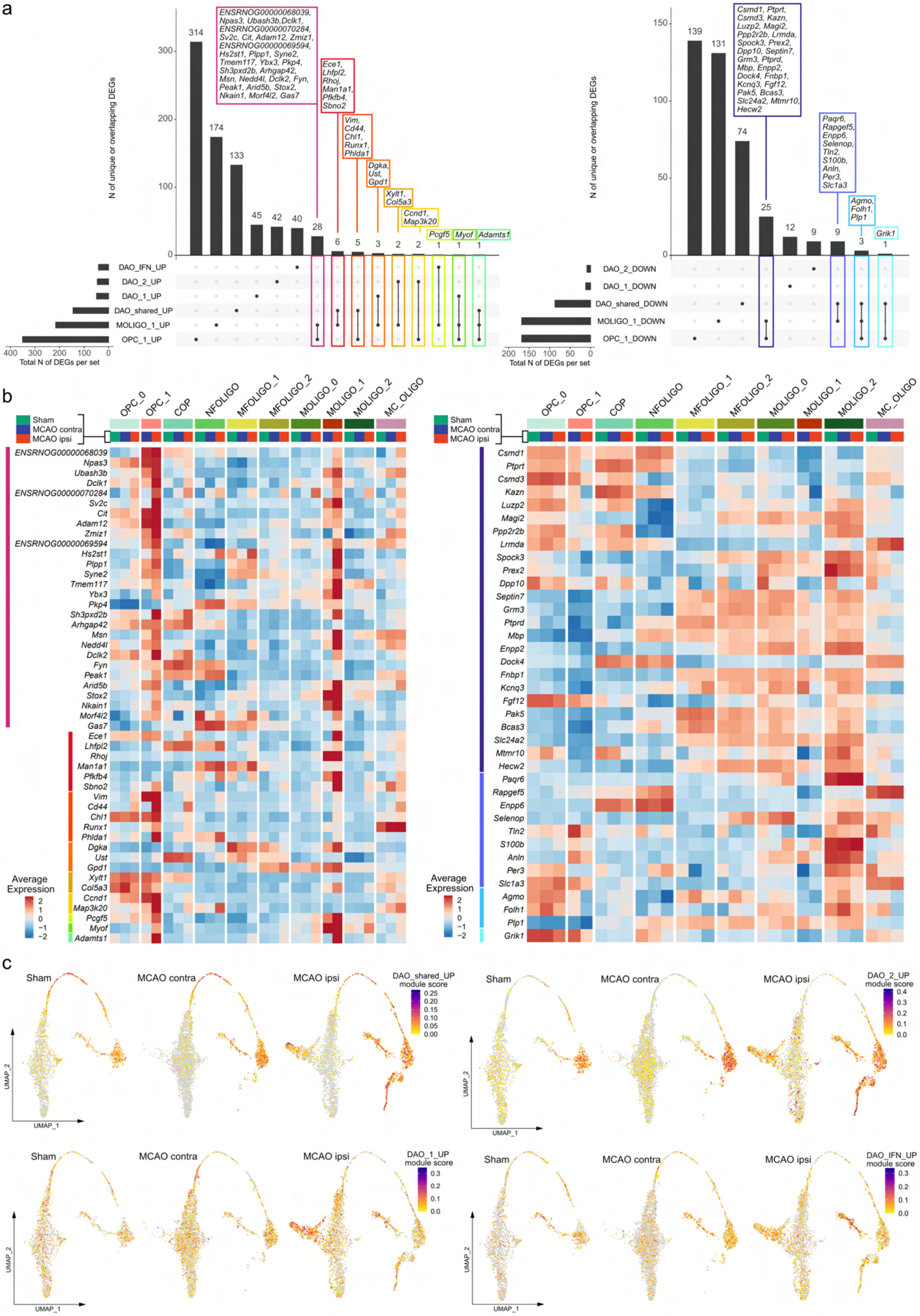


**Supplementary Figure 8. Limited transcriptional overlap between stroke associated oligodendrocyte lineage cells and diseases associated oligodendrocytes (DAO).**
(Legend on next page)

**Supplementary Figure 8. Limited transcriptional overlap between stroke associated oligodendrocyte lineage cells and diseases associated oligodendrocytes (DAO). (a)** UpSet plots depicting the numbers of total, unique and overlapping signature genes of OPC_1, MOLIGO_1 and the DAO signatures: DAO_1 (= immunogenic process associated), DAO_2 (= survival pathway associated) and DAO_IFN (= interferon response associated). Left UpSet Plot compares upregulated, right UpSet plot compares downregulated signature genes. For OPC_1 and MOLIGO_1 all up and downregulated DEGs (Bonferroni-adjusted p-values $< 0.05$ and $|\log2\text{fold change} \geq 0.6|$), derived from the comparison of OPC_1 to OPC_0 and MOLIGO_1 to MOLIGO_2, were included as signature genes, DAO signature gene sets are detailed in Suppl.data.file.1. Overlapping genes are named and color coded. **(b)** Heatmaps depicting the average scaled gene expression of shared signature genes, derived from the comparisons illustrated in **(a)**, split by sub cluster and group. Color codes on the left side of the gene names (y-axes) correspond to color codes in **(a). (c)** Module score feature plots projecting aggregate expression of DAO gene sets on to oligodendrocyte subcluster UMAP plots, split by group.

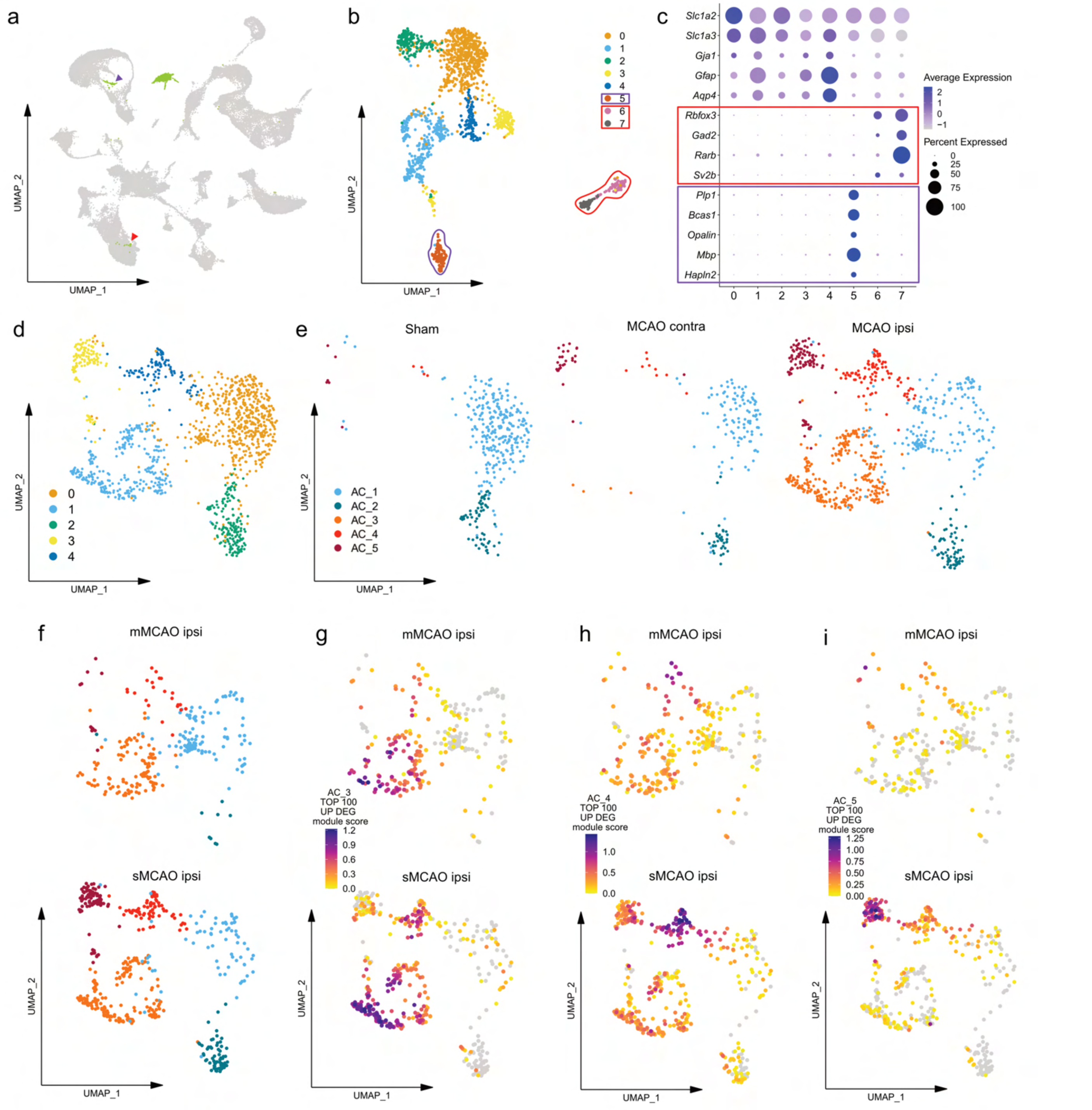


**Supplementary Figure 9. Additional pre-processing steps and between group comparisons pertaining to Figure 4. (a-c)** Removal of neuronal and oligodendrocyte transcript contaminated subclusters. **(a)** Astrocyte cluster (AC) highlighted in the main UMAP plot (green). The arrowhead points to scattered AC nuclei in proximity to oligodendrocyte clusters and the neuronal cluster MSN_2. **(b)** UMAP plot showing non-annotated unsupervised subclustering of AC, with color coded Seurat clusters **(c)** Corresponding dotplot, depicting curated astrocyte, neuronal and oligodendrocyte marker genes. Clusters contaminated with neuronal transcripts (6 and 7) and corresponding neuronal markers are highlighted in red in **b** and **c**. Cluster 5 contaminated with oligodendrocyte transcripts and corresponding oligodendrocyte marker genes are highlighted in violet in **b** and **c**. **(d)** UMAP plot showing non-annotated astrocyte subclusters, after removal of contaminating subclusters. **(e)** UMAP plots depict distribution of nuclei across subclusters, split by treatment group (**e**) and severity (**f**). **(g-i)** The top 100 significantly (Bonferroni-adjusted p-values < 0.05) upregulated DEGs, sorted by descending log2fold changes, derived from the comparison of each reactive astrocyte subcluster (AC_3 to AC_5) to the homeostatic astrocyte subclusters (AC_1, AC_2 pooled) were aggregated to module scores and projected onto subcluster UMAP plots, split by infarction severity (mMCAO = moderate, sMCAO = severe infarctions).

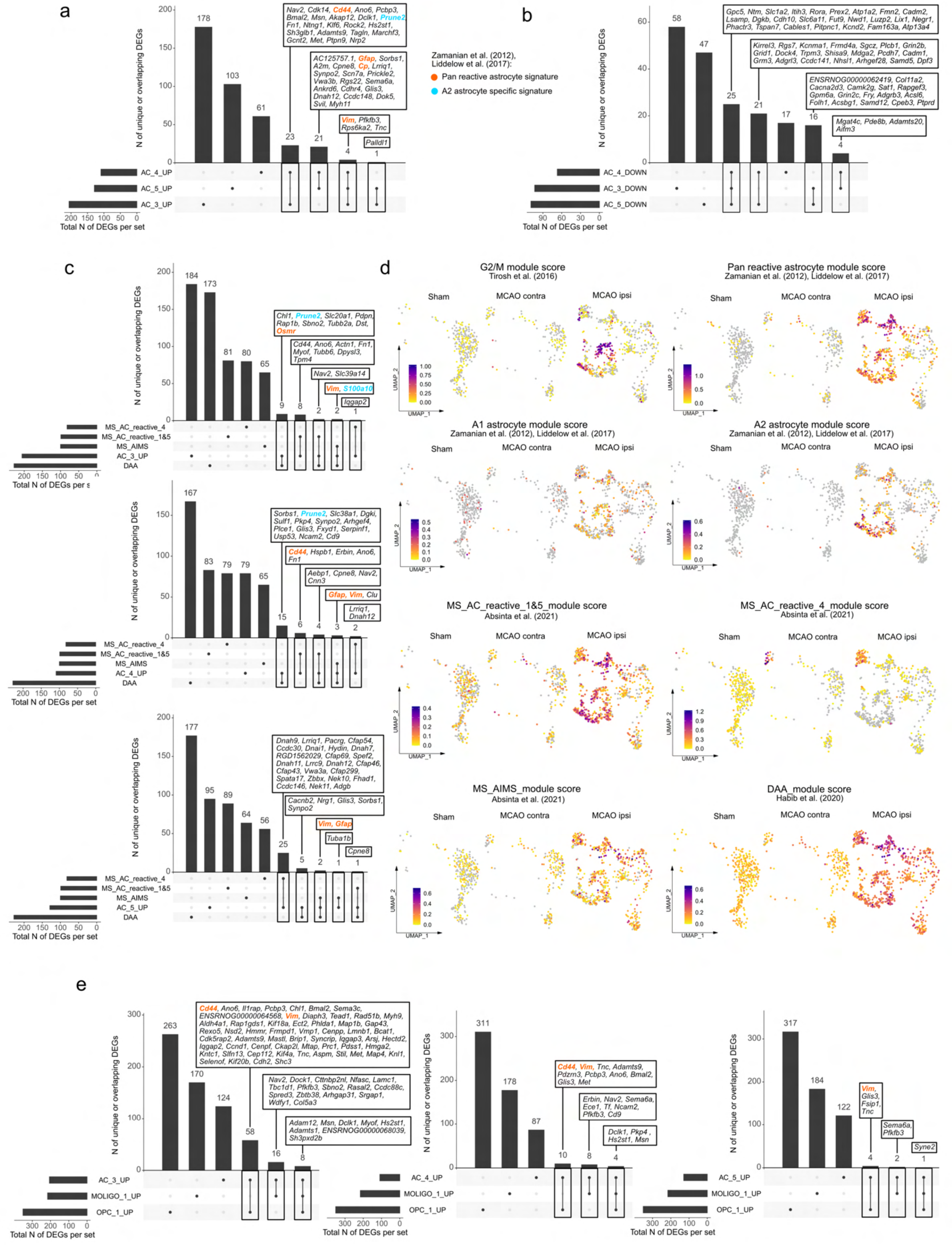


**Supplementary Figure 10. Transcriptional signatures of reactive astrocytes in stroke and other neuropathologies.** (Legend on next page)

**Supplementary Figure 10. Transcriptional signatures of reactive astrocytes in stroke and other neuropathologies. (a,b)** UpSet plots depicting the numbers of total, unique and overlapping signature genes of the stroke associated reactive astrocyte (AC) subclusters (AC_3 to AC_5). All DEGs (Bonferroni-adjusted p-values < 0.05 and |log2fold change ≥ 0.6|) derived from the separate comparison of the reactive astrocyte subclusters AC_3, AC_4, and AC_5 to the homeostatic astrocyte subclusters (AC_1 and AC_2, pooled) were included. **(a)** upregulated, **(b)** downregulated DEGs of AC_3 to AC_5. Genes also included in the pan reactive astrocyte or A2 astrocyte gene sets, are highlighted in orange and cyan, respectively. **(c)** UpSet plots depicting the numbers of total, unique and overlapping signature genes of the stroke associated reactive astrocyte subclusters, as well as reactive astrocyte signatures identified in multiple sclerosis (MS) and neurodegeneration (= disease associated astrocytes (DAA)). Gene sets used in this analysis are detailed in Suppl.data.file.1 **(d)** Module score feature plots projecting aggregate gene expressions of MS reactive astrocyte and DAA signatures onto astrocyte subclustering UMAP plots, split by treatment group. **(e)** UpSet plots depicting the numbers of total, unique and overlapping signature genes in reactive astrocytes and stroke associated oligodendrocyte lineage sub clusters. Up regulated DEGs in AC_3 (left), AC_4 (middle) and AC_5 (right) are compared to upregulated DEGs in OPC_1 and MOLIGO_1.

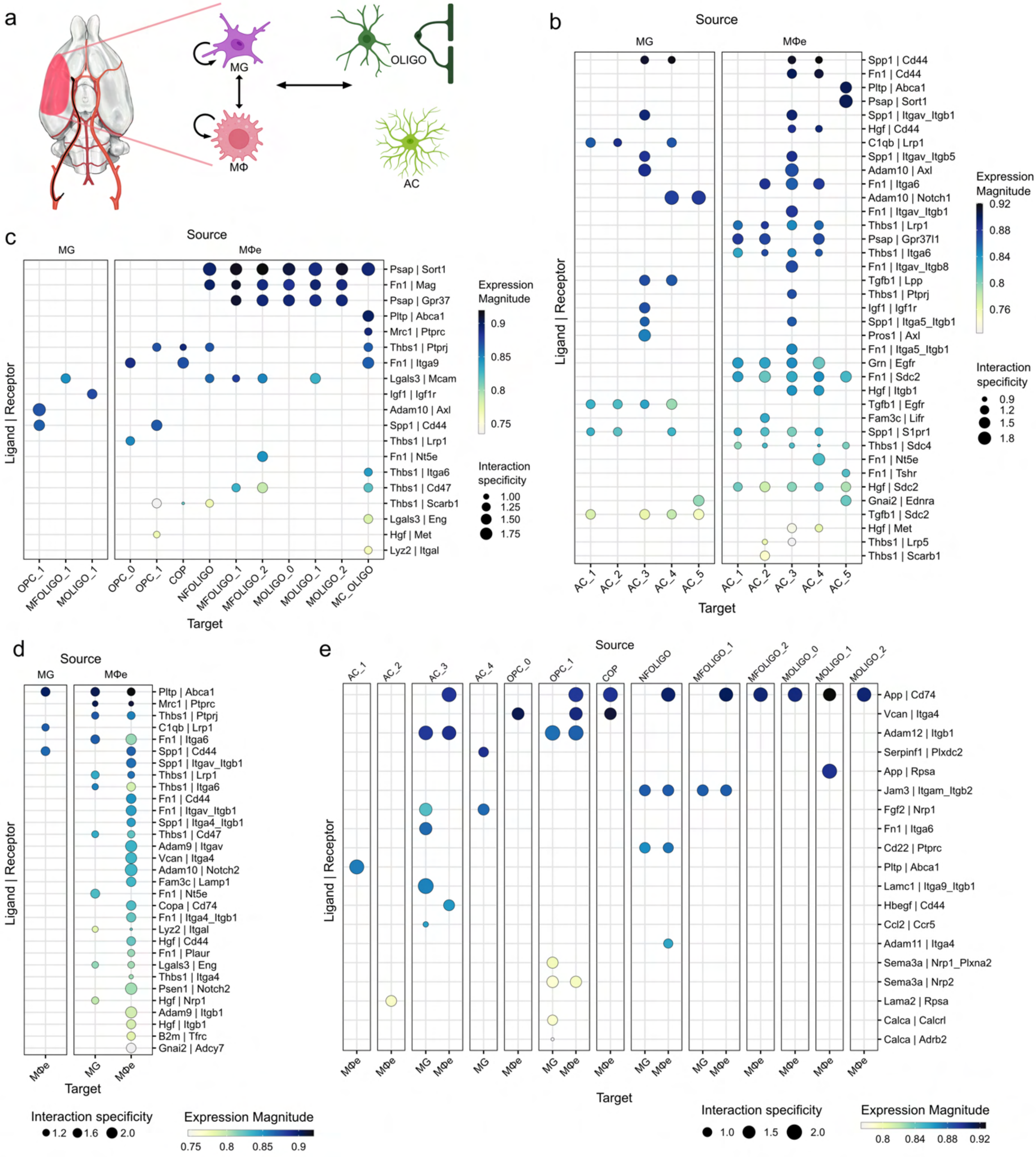


**Supplementary Figure 11. Cell-cell communication analysis infers immuno-glial cross talk within infarcted brain tissue. (a)** Illustration of cell-cell communication events included in this analysis. Illustrated using BioRender.com and SketchBook. **(b-e)** Dotplots depict all significant (aggregate rank scores < 0.05) ligand-receptor (LR) interactions, unique to datasets derived from MCAO ipsi. LR interactions between myeloid cell clusters as sources and astrocyte **(b)** and oligodendrocyte lineage cell clusters **(c)** as targets, between myeloid cell clusters **(d)** and neuroglia clusters as sources and myeloid cell clusters as targets **(e)** are shown. Interaction specificities (= NATMI edge specificity weights) are depicted as dot sizes and LR expression magnitudes (= SingleCellSignalR LR scores) are color coded. Abbreviations: AC: Astrocyte, OLIGO: Oligodendrocyte lineage cells, MG: microglia, MΦ: macrophage, MΦe: macrophage enriched clusters, OPC: oligodendrocyte precursor cell, COP: committed oligodendrocyte precursor, NFOLIGO: newly formed oligodendrocyte, MFOLIGO: myelin forming oligodendrocyte, MOLIGO: mature oligodendrocyte, MC_OLIGO: myeloid cell oligodendrocyte mixed cluster

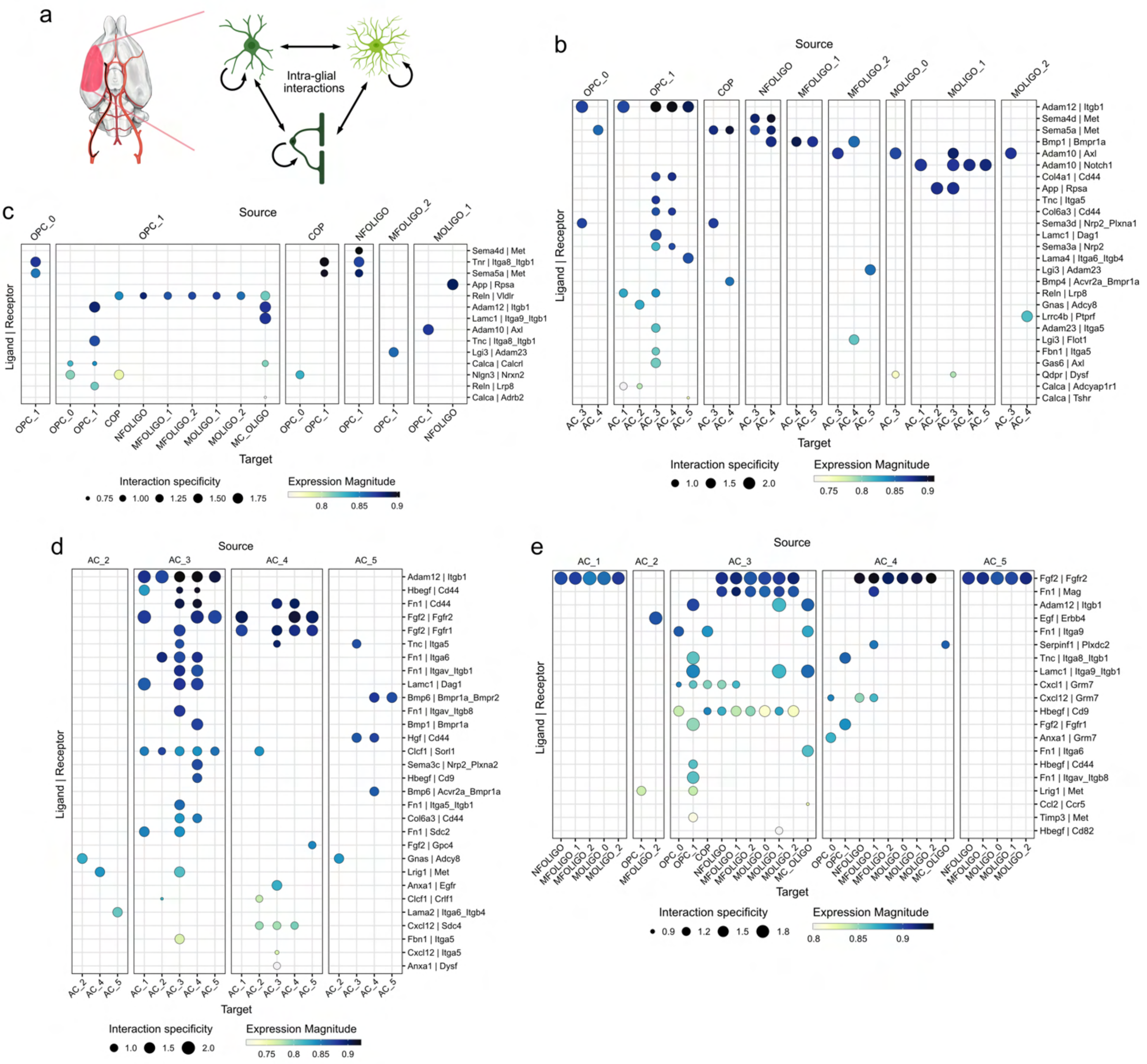


**Supplementary Figure 12. Cell-cell communication analysis infers intra-glial cross talk within infarcted brain tissue. (a)** Illustration of cell-cell communication events included in this analysis. Illustrated using BioRender.com and SketchBook. **(b-e)** Dotplots depict all significant (aggregate rank scores < 0.05) ligand-receptor (LR) interactions, unique to datasets derived from MCAO ipsi. LR interactions between oligodendrocytes as sources and astrocyte **(b)** and oligodendrocyte lineage cell clusters **(c)** as targets, between astrocyte subclusters **(d)** and between astrocyte subclusters as sources and oligodendrocyte subclusters as targets **(e)** are shown. Interaction specificities (= NATMI edge specificity weights) are depicted as dot sizes and LR expression magnitudes (= SingleCellSignalR LR scores) are color coded. Abbreviations: AC: Astrocyte, OLIGO: Oligodendrocyte lineage cells, MG: microglia, MΦ: macrophage, MΦe: macrophage enriched clusters, OPC: oligodendrocyte precursor cell, COP: committed oligodendrocyte precursor, NFOLIGO: newly formed oligodendrocyte, MFOLIGO: myelin forming oligodendrocyte, MOLIGO: mature oligodendrocyte, MC_OLIGO: myeloid cell oligodendrocyte mixed

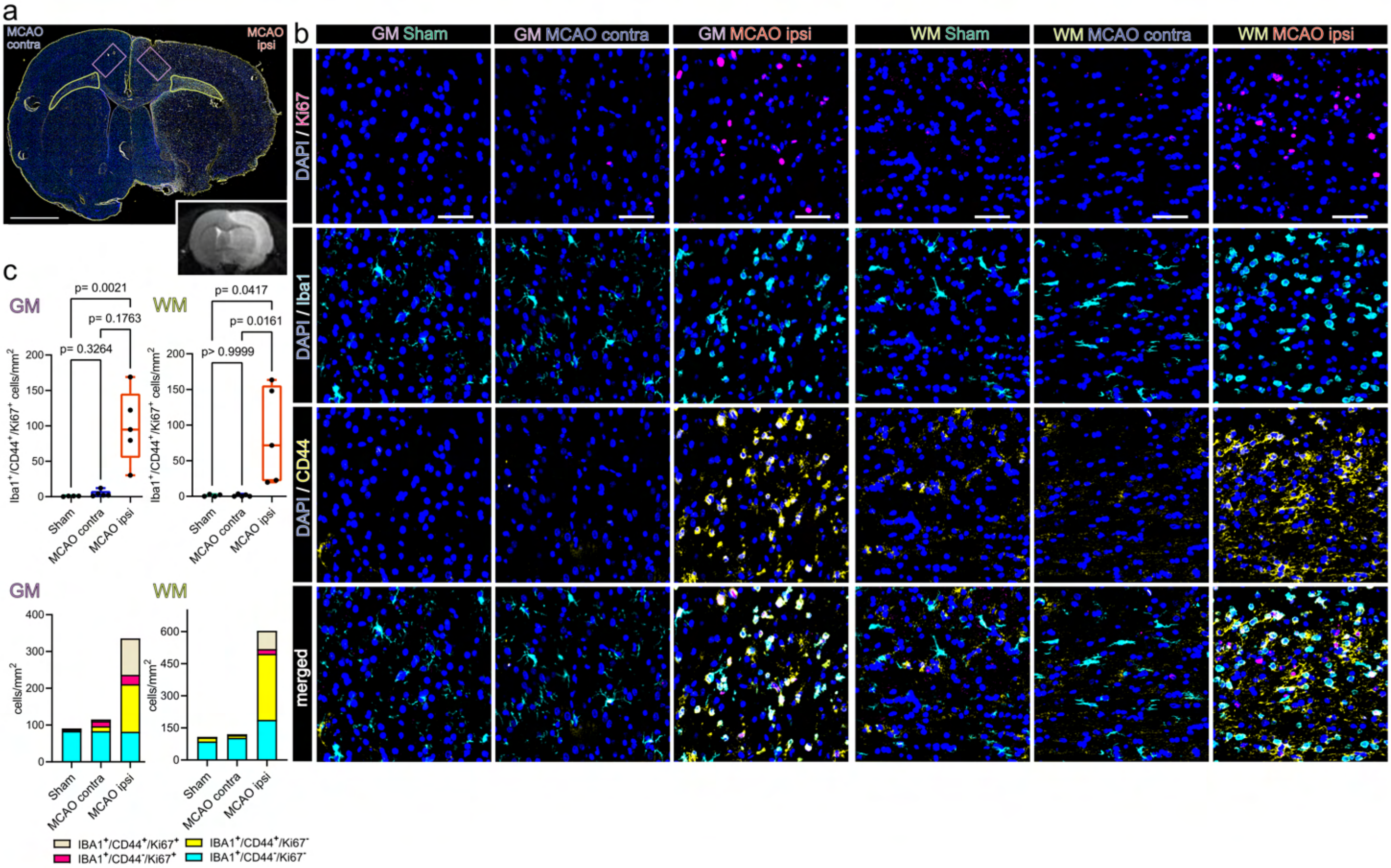


**Supplementary Figure 13. Abundant proliferating, CD44 positive myeloid cells are identified in the perilesional zone surrounding the infarcted area. (a)** Overview of a representative coronal brain section 48h post MCAO, stained for Iba1, CD44 and Ki67. Grey matter ROIs (GM) are highlighted in violet, white matter ROIs (WM) in lime green, lower right inset depicts a corresponding T2 weighted MRI image from the same animal. Bar = 2 mm **(b)** Representative images taken from GM and WM ROIs of Sham, MCAO contra and MCAO ipsi, split by antigen. Ki67 = magenta, Iba1 = cyan, CD44 = yellow, all overlaid with DAPI (nuclei) = blue. Bars = 50 µm. **(c)** Cell counts within GM and WM are presented as box plots for $Iba1^+/CD44^+/Ki67^+$ triple positive cells. Cell counts for $Iba1^+/CD44^+/Ki67^+$, $Iba1^+/CD44^-/Ki67^+$, $Iba1^+/CD44^+/Ki67^-$, $Iba1^+/CD44^-/Ki67^-$ are jointly shown as colored stacked bar plots. Data derived from n = 4-5 animals per group, p values derived from Kruskal-Wallis-H-Tests, followed by Dunn's post hoc comparisons.

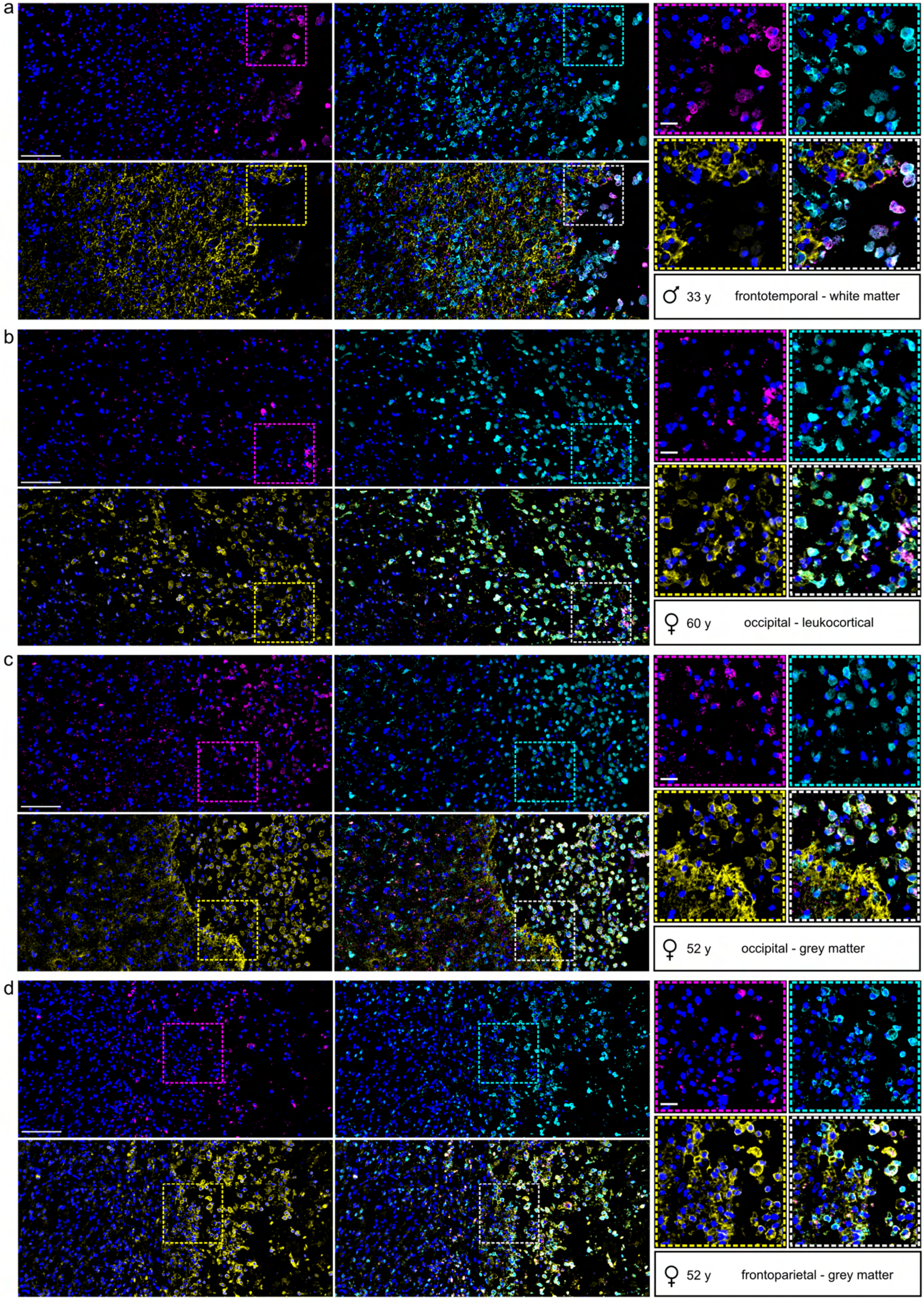


**Supplementary Figure 14. Spatial association of osteopontin positive myeloid cells to CD44 positive cells in human infarcted cerebral tissue.** (Legend on next page)

**Supplementary Figure 14. Spatial association of osteopontin positive myeloid cells to CD44 positive cells in human infarcted cerebral tissue.** Human cerebral, biopsy derived tissue from n = 4 patients, is presented. Iba1, Ki67, OPN and nuclei were visualized using IF staining. **(a - d)** represent individual cases. Sex assigned at birth, age and sampled brain regions are given for each patient. All cases were staged as infarcted tissue in the stage of advanced macrophage resorption and beginning of pseudo cystic cavity formation (Stage II - III). For each case a 400 x 800 µm overview area at the border of macrophage resorption is presented, as well as 150 x 150 µm close up images, split by antigen, clockwise: Ki67 = magenta, Iba1 = cyan, CD44 = yellow, all channels merged = white, all channels were overlaid with DAPI (nuclei) = blue. Scale bars: 100 µm in overview, 20 µm in close ups.

### Supplementary notes – Detailed description of cluster annotation

The overall interpretation of our results regarding cell type specific transcriptional perturbations in response to ischemic stroke is given in the main text. In this supplementary note we describe the performed clustering analyses and the curation of marker genes for cell cluster annotation in detail.

#### Main cell cluster annotation

Unsupervised clustering of the integrated dataset, using the first 23 principal components (PC) and a resolution of 0.4 revealed 32 clusters, which were then annotated to 29 major cell clusters, according to established marker genes (Fig.1b,c). Clusters were first grouped in neuronal and non-neuronal populations using the pan-neuronal markers *Rbfox3*, *Snap25* and *Syt1* [22, 33, 47, 49]. Among the non-neuronal populations, we identified two communities expressing overarching oligodendrocyte lineage markers, such as *Plp1*, canonical markers of oligodendrocyte precursor cells (OPC) and immature oligodendrocytes, such as *Vcan*, *Pdgfra* and *Bcas1* (cluster OLIGO_1) and markers of myelinating and mature oligodendrocytes, such as *Mbp*, *Mog* and *Apod* (cluster OLIGO_2) [31, 53]. We identified one cluster enriched in several canonical pan-astrocyte markers, such as *Slc1a2* and *Slc1a3* [3] and astrocyte associated markers such as *Aqp4* and *Gfap* [37, 53], which we annotated astrocyte cluster (AC). We observed the concomitant enrichment for several ependymal cell marker genes, such as *Tmem212*, *Ccdc162* [53], *Cfap299* [12] and *Dnah11* [37] in one cluster. Notably, the same cluster also faintly co-expressed *Pdgfra*, and several other genes concomitantly expressed in vascular leptomeningeal cell types such as *Dcn*, *Ptgds* and *Aldh1a2* [53]. Thus this cluster was annotated ependymal and mural cell cluster (EP_M_C), due to the possible contribution of both ependymal and mural cell derived transcripts. The expression of canonical endothelial cell markers such as *Pecam1*, *Vwf* and *Flt1* [37, 38] was restricted to one small cluster, which also expressed pericyte associated markers such as *Rgs5* and *Vtn* [8, 38]. Due to the small cluster size this cluster could not be resolved into endothelial, pericyte and mural cell subclusters and was annotated vascular cell cluster (VASC).  Lastly, we identified one immune cell cluster, which was close to exclusively derived from infarcted brain hemispheres. This cluster selectively expressed the broad myeloid cell marker *Ptprc*, the immune cell enriched marker *Arhgap15*, microglia associated transcripts, such as *Hexb*, *Fcrl2* and *Cx3cr1*, shared markers of microglia and macrophages, such as *Aif1* and to a lesser degree macrophage and granulocyte associated marker genes such as *Apoe*, *Lyz2* and *Cd14* [4, 15, 27].

We identified a substantial overlap between the core stroke-associated myeloid cell (SAMC) signature, consisting of the TOP 10 SAMC marker genes: *Spp1, Fabp5, Gpnmb, Ctsb, Ctsl, Lgals3, Lpl, Fth1, Cd63*, and *Ctsd* [4] and the transcriptional signature of the immune cells in our dataset. Notably, lymphoid lineage cluster markers such as the T-cell markers *Cd3e*, *Cd3g*, B-cell markers *Cd19*, *Cd79a*, innate lymphoid cell type marker *Gata3* and NK cell markers *Nkg7* and *Gzma* [4, 27] were virtually absent in our dataset. Details on the subclustering analysis of infarction enriched myeloid cells are given below.

Neuronal clusters were grouped first into excitatory and inhibitory populations, using the established markers *Slc17a7*, *Slc17a6* and *Sv2b* for excitatory, glutamatergic neurons and *Gad1* and *Gad2* for inhibitory, GABAergic neurons [17, 23, 35].

Glutamatergic neurons were first segregated into 2 broad communities, defined by the absence or expression of *Satb2*. Glutamatergic *Satb2* positive (GLU_Satb2+) and negative (GLU_Satb2-) populations were then further characterized by the expression of known markers of topographical identity. In agreement with the observation that *Satb2* is highly enriched in isocortical excitatory neurons [18, 37], the marker genes of the *Satb2* positive populations GLU_Satb2+_1 to 9, largely overlapped with established transcriptional signatures associated to specific cortical layers (Fig. 1c).

GLU_Satb2+_1 exhibited concomitant enrichment for the L2 associated marker *Otof* [51], the recently identified L2/3 specific marker *Ccbe1* [6], as well as the L2/3 associated markers *Rasgrf2, Glis3, Nectin3, Cux1* and *Cux2* [22, 23, 45, 51, 54]. *Cux1* and *Cux2* were also enriched in GLU_Satb2+_2 and GLU_Satb2+_8. Notably, *Cux1* and *2* are also expressed in deeper cortical layers, for example adjacent to the Claustrum, or layer 4 cortical neurons [6, 51]. The established cortical layer 4-5 marker *Rorb* [23, 37], as well as the *Rorb* target *Thsd7a* [7] were concomitantly enriched in GLU_Satb2+_2 and GLU_Satb2+_3. GLU_Satb2+_3 also expressed the cortical layer 4 to 5 associated marker *Il1rapl2* [23], while the layer 5 and 6 associated markers *Tox* and *Grik3*, [22, 23, 37], where co-expressed to varying degrees from GLU_Satb2+_3 to GLU_Satb2+_7. GLU_Satb2+_4 to 7 expressed *Bcl11b*, which was shown to be highly expressed in layer 5 cortical neurons [26]. Correspondingly, GLU_Satb2+_4 concomitantly expressed the layer 5 associated marker *Parm1* [38] and *Serpine2*, which was previously used as markers of cortical Layer 5b identity in the adult rodent motor cortex [36].

GLU_Satb2+_5 to 7 prominently expressed *Tle4*, which recent evidence suggested as a specific marker of layer 6 corticothalamic projection neurons [46]. GLU_Satb2+_5 and 6 also expressed the corticothalamic projection neuron associated marker *Thsd7b* [51]. GLU_Satb2+_6 was strongly enriched for *Foxp2*, which was associated to cortical layers 5 and 6 [23, 44], as well as the layer 6 associated marker *Syt 6* [37, 54]. Of note, the *Tle4*, *Foxp2* double positive cluster GLU_Satb2+_7, strongly expressed *Vwc2l*, a marker of L5/6 near-projecting neurons [51]. This cluster however also expressed the lateral cortex layer 6 marker *Col24a1* [53] and the cortical layer 5 and 6 associated marker *Tshz2* [44], whereas GLU_Satb2+_8 most prominently expressed the L6 and claustrum associated markers *Synpr* [9, 54] and *Nr4a2* [37, 54] , corroborated by the prominent and selective expression of the claustrum enriched gene *Gng2* (Fig. 1c). GLU_Satb2+_7 and 8 further shared the concomitant expression of *Grin3a* and the layer 5 and 6 associated marker *Htr2c* [22].

GLU_Satb2+_9 shared several L5 and L6 associated features with GLU_Satb2+_3, but distinctly expressed several allocortex associated marker genes, such as *Rspo2* and *Rxfp1*, which are known to be enriched in subcortical structures such as the basolateral amygdala [40], the hippocampal formation, or various diencephalic structures [30]. GLU_Satb2+_9 further exhibited enrichments for *Sema3e* and *Adamts3*.

GLU_Satb2-_1 and 2 both coexpressed *Abi3bp* and *Ndst4*, which have been associated to glutamatergic neurons in anterior olfactory areas [53]. In contrast to its neighbouring cluster, GLU_Satb2-_1 strongly and specifically expressed *Ankfn1*, which was shown to be enriched in the superchiasmatic nucleus [29]. This cluster also prominently expressed *Ntng1* which was shown to be particularly enriched in hypothalamic and thalamic areas [50].

GLU_Satb2-_3 showed a strong enrichment for the prosubiculum associated marker *Arhgap6* and expressed *Rerg* and *Pkp2*, which are enriched in the hippocampal formation [51]. Congruently, this cluster was also enriched for *Rasgrf2* and *Rfx3*, which are particularly densely expressed in upper layer cortical and hippocampal pyramidal cells (Fig. 1c).

We identified one cluster (CHOL_IN) co-expressing *Chat*, *Lhx8*, *Slc17a8* and *Slc5a7*, matching the transcriptional signature of cholinergic interneurons [53].

GABAergic populations were segregated into clusters matching known interneuron (GABA_IN) and GABAergic medium spiny neuron identities (GABA_MSN).

In accordance with previously established annotation strategies [17, 44, 51] we first broadly separated the GABAergic neuronal clusters into *Adarb2* positive (GABA_IN_Adarb2+), thus likely caudal ganglionic eminence (CGE) derived and *Adarb2* negative (GABA_IN_Adarb2-), thus likely medial ganglionic eminence (MGE) derived inhibitory interneuron communities.

Congruently, GABA _Adarb2+_1 selectively expressed *Vip*, [17, 23, 44, 51] and *Cnr*, which are known to be enriched in several *Adarb2* positive corticohippocampal interneurons [53].

GABA _Adarb2+_2 was selectively enriched for the CGE interneuron associated markers *Jam2*, and *Lamp5* [51], and expressed *Sv2c* and *Reln*, previously used to denote several cortical interneuron subsets [23, 47], this cluster also strongly expressed *Cyct* and *Sema5a*.

In agreement with their presumed MGE origin the Adarb2 negative clusters GABA_Adarb2-_1-5 were all *Sox6* positive as previously described [51] and GABA _Adarb2-_1,2 and 4 , further expressed *Satb1*, a previously characterized driver of the terminal differentiation of MGE derived interneurons. As expected these *Adarb2*-, *Sox6*+ populations were also *Lhx6* positive albeit faintly in our dataset.

The canonical MGE interneuron subset marker *Sst* [17, 44, 51] was expressed robustly in GABA _Adarb2-_1. Notably, The distinct concomitant expression of *Sst*, *Nos1*, *Npy*, *Tacr1* and *Chodl*, observed in this cluster matches the transcriptional profile of cortico-hippocamapal, long-range projection interneurons [53]. However, the coexpression of *Sst*, and *Npy* has also been described in distinct subsets of striatal [34] and hypothalamic [43] GABAergic interneurons, brain regions which were also sampled in our study.

Only few cells in the neighbouring cluster GABA _Adarb2-_2 were *Sst* positive, however the concomitant expression of *Grik3*, *Elfn1* and *Lypd6* observed in this cluster was reminiscent of the transcriptional profile of *Sst* positive hippocamposeptal interneurons [53]. Of note, the coexpression of *Grik3* and *Elfn1* was also observed in GABAergic interneurons of the globus pallidus externus [38].

Several markers of previously described MGE interneuron subsets, such as *Etv1*, *Pde3a* [51], and *Eya1* [44], were selectively enriched in GABA _Adarb2-_3, which was also strongly *Dpf3* positive.

Interestingly, the axon guidance and progenitor cell proliferation master regulator *Slit 2* [13] was selectively enriched in GABA_Adarb2-_4. This *Pvalb* positive interneuron population was also enriched in *Eya4*, as well as *Rspo2* and *Esrrg*, known to be enriched in the amygdala and thalamus, respectively [53].

GABA_Adarb2-_5 interneurons distinctly coexpressed *Col5a2*, *Hs3st2*, *Uncb5* and *Vipr2*. *Vipr2* has been previously suggested as a marker of a distinct subset of *Pvalb* interneurons [44, 51]. However, in our dataset the *Vipr2* positive cluster only contained few *Pvalb* positive cells. Likewise, *Unc5*b, which was suggested as a marker of *Pvalb*+ chandelier cells [17], was in our dataset selectively expressed in cluster GABA_Adarb2-_5.

Strikingly, two GABAergic communities distinctly coexpressed *Rgs9*, *Rarb* and *Gng7*, markers which are largely restricted to the basal ganglia and the cortical sub plate [25, 37]. These clusters matched previously validated transcriptional profiles of striatal medium spiny projection neurons and were annotated GABAergic medium spiny neuron clusters 1 and 2 (GABA_MSN_1 and _2). GABA_MSN_1 selectively expressed *Tshz1*, which is known to be densely expressed in olfactory areas, the thalamus and the basal ganglia, while in the later it has been established as a marker of striosomal direct pathway MSNs [38, 48]. This cluster additionally robustly coexpressed, *Adarb2*, *Foxp2*, and *Olfm3* and might thus harbour a recently described, distinct *Foxp2/Olfm3* double positive striatal subpopulation [24], further corroborated by the selective expression of *Otof*, which among GABAergic neurons has been suggested as a marker of eccentric MSNs [38]. *Rgs9, Rarb, Gng7* and the basal ganglia associated orphan GPCR gene *Gpr149* [41] were expressed more prominently in GABA_MSN_2, which also expressed markers of canonical striatal medium spiny projection neurons such as *Ppp1r1b* and *Drd2* [38, 53].

In comparison, the remaining GABAergic community, lacked specific markers and did not match to known transcriptional interneuron and MSN identities, and was thus termed ambiguous GABAergic cell cluster (GABA_Amb). Most notably, this cluster distinctly coexpressed *Slc17a6*, which was found to be expressed in Gad1/2 positive cells of deep grey matter structures such as the hypothalamus [33, 43] and the cortical sub plate enriched gene *Glra3*. It is thus plausible that several non-cortical interneuron populations might have contributed to this cluster.

**Subclustering of infarction enriched myeloid cells**

Unsupervised subclustering of MCAO ipsi derived MCs (n= 2646 nuclei) using the first 10 PCs and a resolution of 0.4 corroborated the myeloid cell identity of this cluster revealing 6 subclusters (Suppl.Fig.3c-e). Two clusters (MG_0 and MG_1) where predominantly enriched in pan microglia markers such as *C1qa* and *Fcrls*, expressed the homeostatic microglia markers *Cx3cr1*, *P2ry12* and *Tmem119* [15, 32] to varying degrees, but where almost devoid of macrophage associated transcripts. Of note we observed a higher average expression of homeostatic microglia markers in the more M0-like cluster MG_0 than in MG_1. Three clusters, only sparsely expressed microglia marker genes, however where enriched in the macrophage associated transcripts *Apoe* and *Lyz2*, other macrophage associated transcripts such as *Cd14*, *Ccr2*, *Fn1* and *Cybb* where expressed to variable degrees in these cluster [4, 27]. We thus reasoned that these cluster are primarily derived from macrophage nuclei. However, previous work denoted an extensive overlap of the transcriptional signatures of microglia and CNS macrophages and a loss of microglia marker signatures under several pathological conditions, as well as acquisition of macrophage associated markers such as *Apoe* and *Lyz2* im microglia [4, 27, 32, 42]. Therefore, these clusters were annotated macrophage enriched clusters MΦe_1 to 3, conceding that a contribution of microglia to these subclusters cannot be excluded. The remaining cluster was strongly enriched for the dendritic cell associated marker *Flt3* [27] and accordingly most robustly expressed *Cd74* and several rat orthologues of the human major histocompatibility class II complex such as *RT1-Bb* and *RT1-Db1*. This cluster was therefore annotated dendritic cell cluster (DC).

**Subclustering of oligodendrocyte lineage cells**

Joint unsupervised sub clustering of OLIGO_1 and OLIGO_2, using the first 13 PCs and a clustering resolution of 0.3, revealed 12 sub clusters (Fig.2). Two sub clusters (total n= 639 nuclei) only faintly expressed canonical oligodendrocyte lineage markers but were enriched for *Rbfox3, Rarb*, *Gad2* and *Foxp2*, reminiscent of MSN neurons. Due to this biologically implausible marker combination these clusters were deemed to be clustering artefacts and removed from downstream analyses (Suppl.Fig.6a-d). Thereafter, the dataset was reclustered, at the above described parameters. The remaining 10240 nuclei split into 10 sub clusters. 3 populations were enriched for the shared OPC and committed oligodendrocyte precursor (COP) associated markers *Sox6* and *Vcan* [16, 19, 31], of which 2 populations (OPC_0, OPC_1) also expressed canonical OPC markers such as *Pcdh15*, *Ptprz1*, *Pdgfra*, *Cspg4*, or *C1ql1* [16, 31, 53]. Branching of OPC_0 we identified one cluster with a residual *Pcdh15* and *Ptprz11* expression and strong enrichment for the canonical COP markers *Bmp4, Neu4* and *Tnr*, COP migration associated genes *Tns3* and *Fyn* and the immature oligodendrocyte marker *Bcas1* [11, 31, 53]. The COP cluster also strongly expressed genes, which have been shown to be upregulated upon the transition of COPs to newly formed oligodendrocytes (NFOLIGO), such as *Tcf7l2* or *Itpr2* [31]. Next to the COP cluster we identified one cluster positive for these genes, concomitantly expressing the *bona fide* NFOLIGO markers *Rras2* and *Cnksr3* [53]. These markers where residually expressed in the adjacent myelin forming oligodendrocyte cluster 1 (MFOLIGO_1). Both myelin forming oligodendrocyte clusters (MFOLIGO_1 and MFOLIGO_2) exhibited a robust concomitant enrichment for the markers of myelin formation *Opalin, Serinc5, Mbp , Mobp, Mal, Mog, Plp1*, [16, 19, 31, 38, 53], as well as the early stage myelination associated marker *Tspan2* [5]. Importantly, these clusters distinctly lacked the expression of the late stage oligodendrocyte differentiation gene *Hapln2* [16, 37, 53] and only faintly expressed other mature oligodendrocyte associated genes, such as *Dock5* and *Apod* [16, 31]. The clusters co-expressing the established oligodendrocyte maturity markers *Dock5, Hapln2* and *Apod*, were termed mature oligodendrocyte clusters (MOLIGO_1 to 3).

We identified one sub cluster coexpressing oligodendrocyte transcripts and immune cell (e.g. *Ptprc*, or *Fcrl2*) and immune process (e.g. *Cfh, Anxa3, Lyn*) associated genes. As detailed in the main text, previous research indicated the possibility of oligodendrocyte transcript phagocytosis by myeloid cells and enrichment of these transcripts within the nuclear compartment, in pathological conditions such as in MS lesions [39]. Congruently, particularly myeloid cell markers (e.g. *Ptprc, Fcrl2*), SAMC associated genes (e.g. *Gpnmb, Spp1*) and the phagocytosis related gene *Anxa3* [20], were derived from infarcted, but not Sham group brain tissue (Suppl.Fig.6e). However previous research also described the occurrence of immune process related genes in bona fide OPCs and oligodendrocytes termed immune oligodendroglia [10, 19, 21]. We thus concede that immune oligodendroglia transcript might have also contributed to this cluster. However, due to the obvious contribution of myeloid cells we opted to annotate this subcluster as a myeloid cell oligodendrocyte mixed cluster (MC_OLIGO).

**Subclustering of astrocytes**

Unsupervised sub clustering of astrocytes, using the first 14 principal components (PCs) and a clustering resolution of 0.3, revealed 8 sub clusters. Similar to the results of the oligodendrocyte subclustering analysis we observed two sub clusters, with evident neuronal transcript contamination (e.g. *Rbfox3*). We also observed one cluster which only weakly expressed pan-astrocyte markers, such as *Slc1a2*, *Slc1a3*, or *Gja1* or markers of reactive astrocytes, such as *Gfap* or *Aqp4*. Recent research provided evidence for the reprograming of mature oligodendrocytes into astrocytes, via a Plp+/GFAP+ "AO"-stage during acute brain injury [2]. However, within the astrocyte subclustering in our dataset, the subcluster concomitantly expressing astrocyte and oligodendrocyte transcripts lacked reactive astrocyte marker genes, was conserved across all datasets and not associated to infarcted hemispheres. Furthermore, DEG analyses comparing the transcriptional profiles within this cluster between Sham hemispheres, infarcted hemispheres and hemispheres contralateral to infarction did not reveal any DEGs. Because we thereby excluded that this cluster harbors a stroke specific gene expressional signature in line with the AO-stage, we treated this cluster as a technical artefact and did not consider it in further calculations. In total we removed 225 nuclei from downstream analyses and the remaining 1233 nuclei were reclustered at the above described parameters, revealing 5 filtered sub clusters.

Astrocyte subclusters could be broadly split into homeostatic astrocyte gene enriched (e.g. *Gpc5, Kirrel3, Cdh10, Trpm3*) [1, 14, 39] sub clusters (AC_1 and AC_2) and reactive astrocyte clusters (AC_3 to AC_5), expressing reactive astrocyte associated genes, such as *Gfap, Vim, Osmr, Cd44,* or *Cp*) [28, 52]. AC_2 differed from AC_1 in the expression of *Slc7a10*, which is more enriched in astrocytes of the olfactory area and *Agt* and *Slc6a11* which are restricted to non telencephalic astrocyte subsets [53].
